## Supplementary material for "The *Drosophila drop-dead* gene is required for eggshell integrity": File S1: neutral red staining protocol

**Neutral red staining of Drosophila eggs**

Reagents/equipment:
Grape juice agar plates for egg collection
Yeast paste
12-well plate
5 mg/ml neutral red solution, filtered (will last for months at room temp)
bleach
wire mesh egg basket
curved forceps
1 mL pipetteman and tips
MilliQ water
beaker for liquid waste
timer

Protocol:

1. Collect eggs on a grape juice agar plate with yeast paste. The length of the collection will depend on the experiment, from a couple of hours to overnight, depending on how synchronized you need the eggs to be.
2. Prepare the 12-well plate by putting filling wells with 2 mL each of the following solutions, in order: water, 50% bleach, water, water, neutral red, water, water
3. Gather laid eggs in one section of the plate
4. Put the basket in the first well of water and transfer a known number eggs into the basket using fine forceps (don't actually try and grab eggs with the forceps as you will damage them). The maximum number of eggs I've ever done in one assay is 50.
5. Move the basket into the bleach well for three minutes. Agitate the basket several times during the three minutes, and try to knock the eggs off of the surface by dropping drops of bleach onto them.
6. After three minutes, hold the basket over the waste beaker and wash several times with pipettes full of water. Then move the basket to the next well of water, mix briefly, and move to the next well of water.
7. Under the dissecting scope, count the eggs in the basket to determine how many lysed during the bleach treatment.
8. Move the basket to the neutral red well for 10 minutes. Knock the eggs off of the surface as in step five.
9. After 10 minutes, wash the basket over the waste beaker as in step six, then transfer sequentially to the last two wells of water.
10. Under the microscope, count the eggs and score into the following categories: not dechorionated, unstained and intact, stained and intact, stained and leaking.
11. Analyze pooled data through a series of binary chi-square contingency tests: i.e. did groups differ in the number of eggs that lysed during bleaching, the number of surviving

eggs that were dechorionated, the number of dechorionated eggs that were permeable to dye.
