## Supplementary material for "The *Drosophila drop-dead* gene is required for eggshell integrity": Raw images of all Western blots used for figures

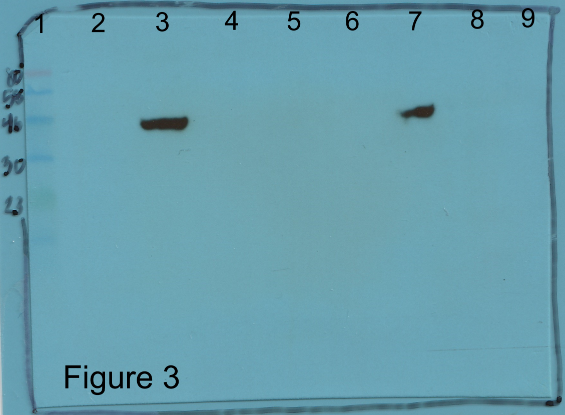


Figure 3

**Blot for Figure 3.**

Lane 1: MW markers

Lane 2: *drd*$^{1}$/*drd*$^{1}$, stage 10 egg chamber

Lane 3: *drd*$^{1}$/*drd*$^{1}$, stage 14 egg chamber

Lane 4: *drd*$^{1}$/*drd*$^{1}$, laid eggs 0-3 hours after oviposition

Lane 5: *drd*$^{1}$/*drd*$^{1}$, laid eggs 0-6 hours after oviposition

Lane 6: *drd*$^{1}$/*FM7c*, stage 10 egg chamber

Lane 7: *drd*$^{1}$/*FM7c*, stage 14 egg chamber

Lane 8: *drd*$^{1}$/*FM7c*, laid eggs 0-3 hours after oviposition

Lane 9: *drd*$^{1}$/*FM7c*, laid eggs 0-6 hours after oviposition

4 egg chambers or eggs per lane. Anti-Cp36, 1:5000.

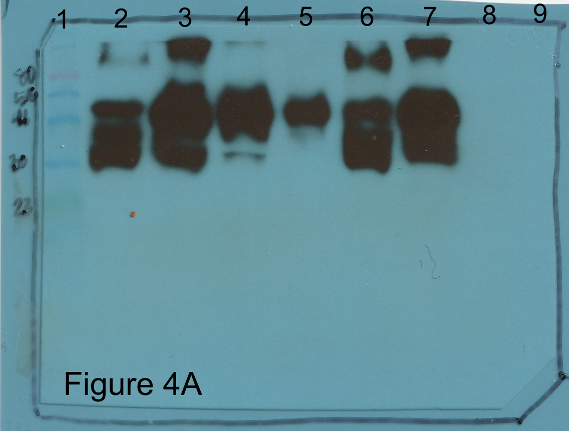


Figure 4A

**Blot for Figure 4A**

Lane 1: MW markers

Lane 2: $drd^{1}/drd^{1}$, stage 10 egg chamber

Lane 3: $drd^{1}/drd^{1}$, stage 14 egg chamber

Lane 4: $drd^{1}/drd^{1}$, laid eggs 0-3 hours after oviposition

Lane 5: $drd^{1}/drd^{1}$, laid eggs 0-6 hours after oviposition

Lane 6: $drd^{1}$/*FM7c*, stage 10 egg chamber

Lane 7: $drd^{1}$/*FM7c*, stage 14 egg chamber

Lane 8: $drd^{1}$/*FM7c*, laid eggs 0-3 hours after oviposition

Lane 9: $drd^{1}$/*FM7c*, laid eggs 0-6 hours after oviposition

4 egg chambers or eggs per lane. Anti- Vm26Ab, 1:10,000.

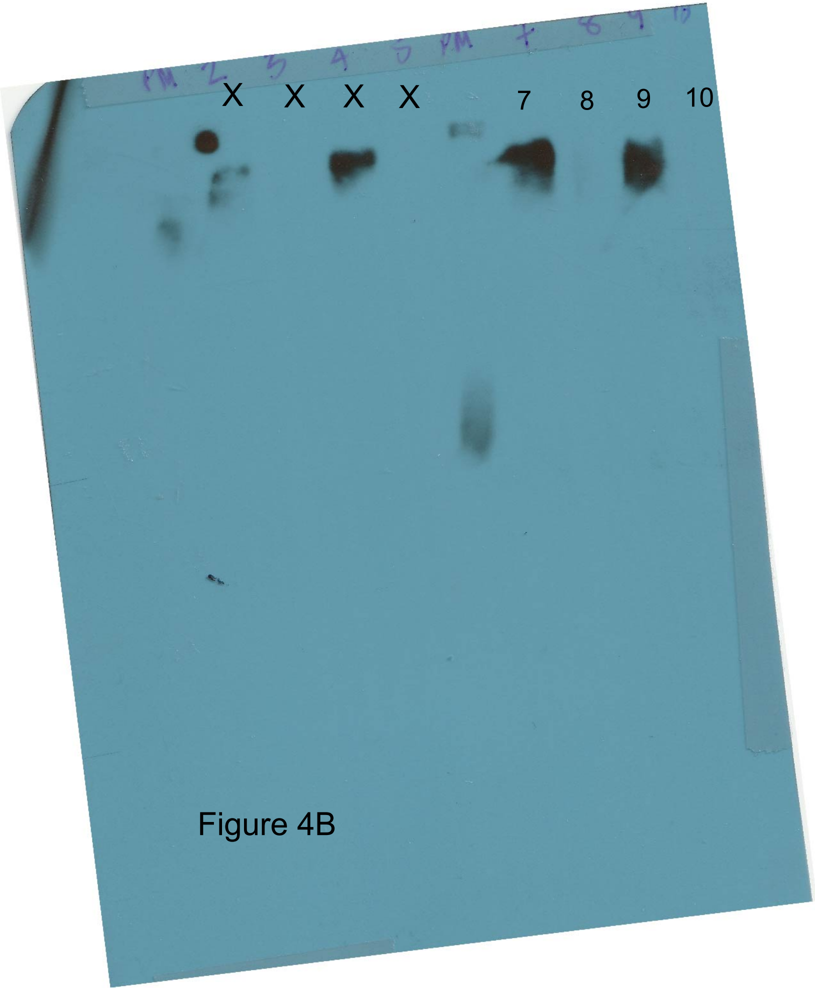


Figure 4B

**Blot for Figure 4B**

Lane 7: *drd*$^{1}$/*FM7c*, stage 10 egg chamber

Lane 8: *drd*$^{1}$/*FM7c*, stage 14 egg chamber

Lane 9: *drd*$^{1}$/*drd*$^{1}$, stage 10 egg chamber

Lane 10: *drd*$^{1}$/*drd*$^{1}$, stage 14 egg chamber

2 egg chambers per lane, no reducing agents added. Anti- Vm26Ab, 1:10,000.

Note that lanes 2-5 are the same samples, same antibody, 1:25,000 dilution.

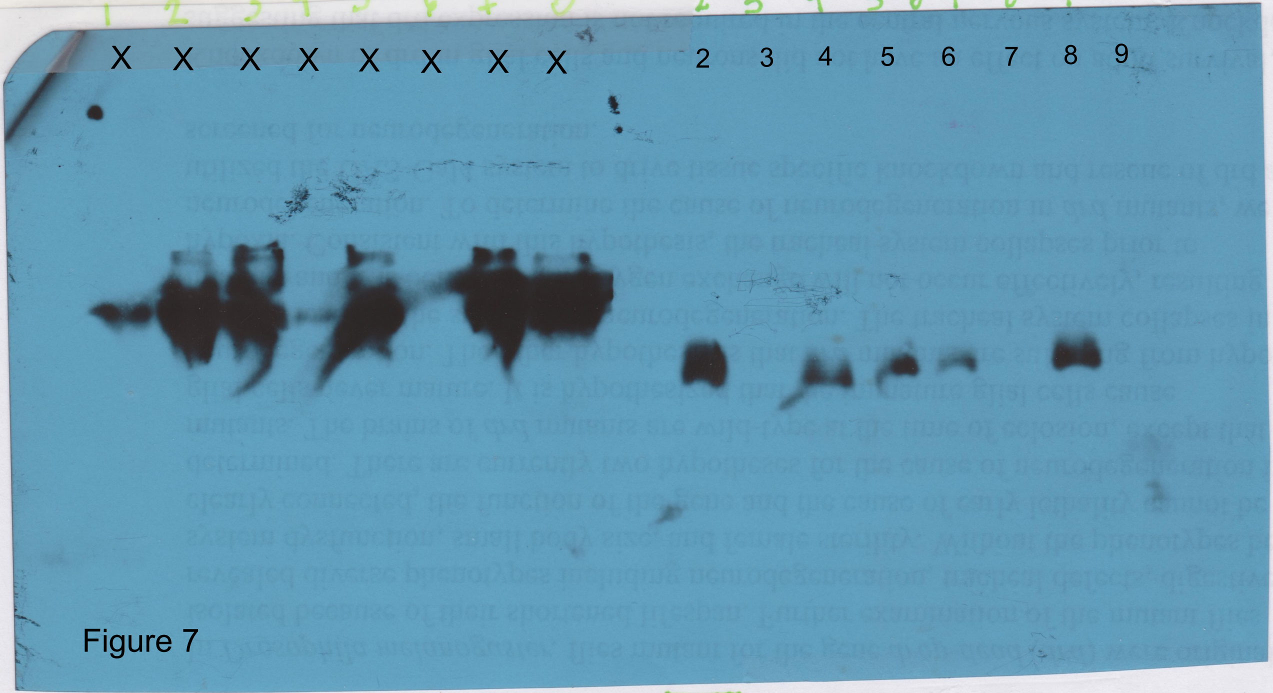


Figure 7

**Blot for Figure 7**

Lane 2: *w; CY2-GAL4/CyO*, stage 14 egg chambers

Lane 3: *w; CY2-GAL4/CyO*, laid eggs

Lane 4: *w; UAS-Dcr-2 drd*$^{GD3367}$*/CY2-GAL4*, stage 14 egg chambers

Lane 5: *w; UAS-Dcr-2 drd*$^{GD3367}$*/CY2-GAL4*, laid eggs

Lane 6: *w; CyO/+; T155-GAL4/+*, stage 14 egg chambers

Lane 7: *w; CyO/+; T155-GAL4/+*, laid eggs

Lane 8: *w; UAS-Dcr-2 drd*$^{GD3367}$*/+; T155-GAL4/+*, stage 14 egg chambers

Lane 9: *w; UAS-Dcr-2 drd*$^{GD3367}$*/+; T155-GAL4/+*, laid eggs

2 egg chambers or eggs per lane. Anti- Vm26Ab, 1:25,000.
