## Supplementary material for "The *Drosophila drop-dead* gene is required for eggshell integrity": Figure S1

**wild-type**

**TAATT**GTAGGTATTCGCCCAGGTCTAGCATTTATAGAGAATTGTATTTTTTGTTGTTTTTTATTTGCTCAACGCAG**CTCGA**

***drd*[1]**

**TAATT**GTAGGTATTCGCCCAGGTCTAGCATTTATAGAGAATTGTATTTTTTGTTGTTTTTTATT<u>A</u>G**CTCAACGCAGCTCGA**
