## Supplementary figures and images for "The *Drosophila drop-dead* gene is required for eggshell integrity"

### Figure S2

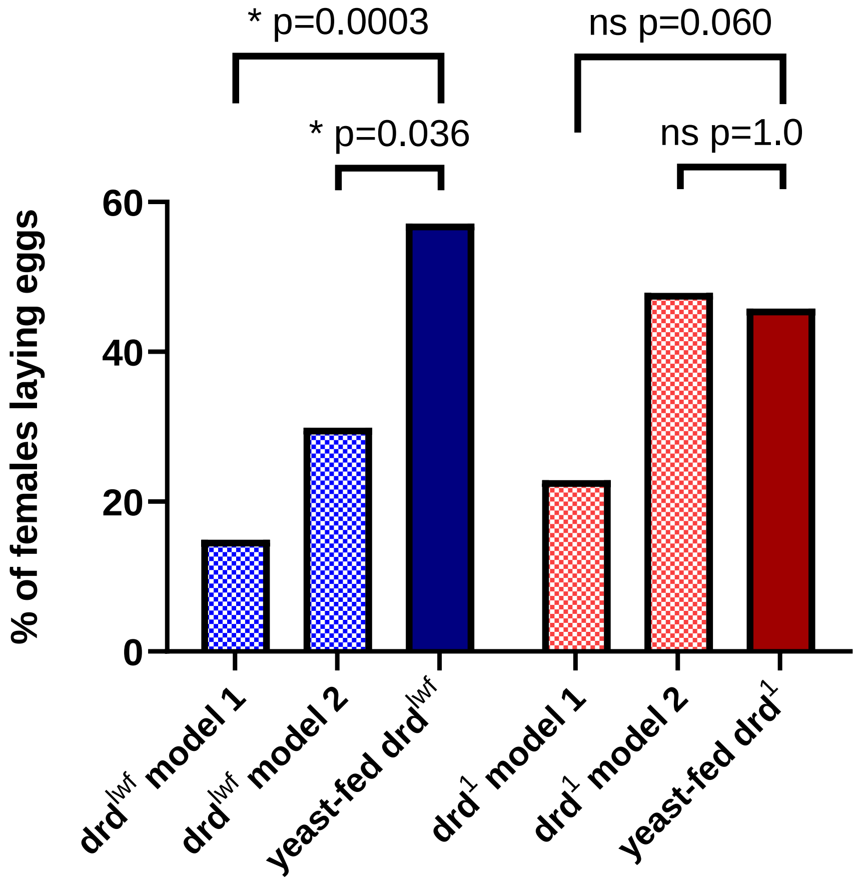

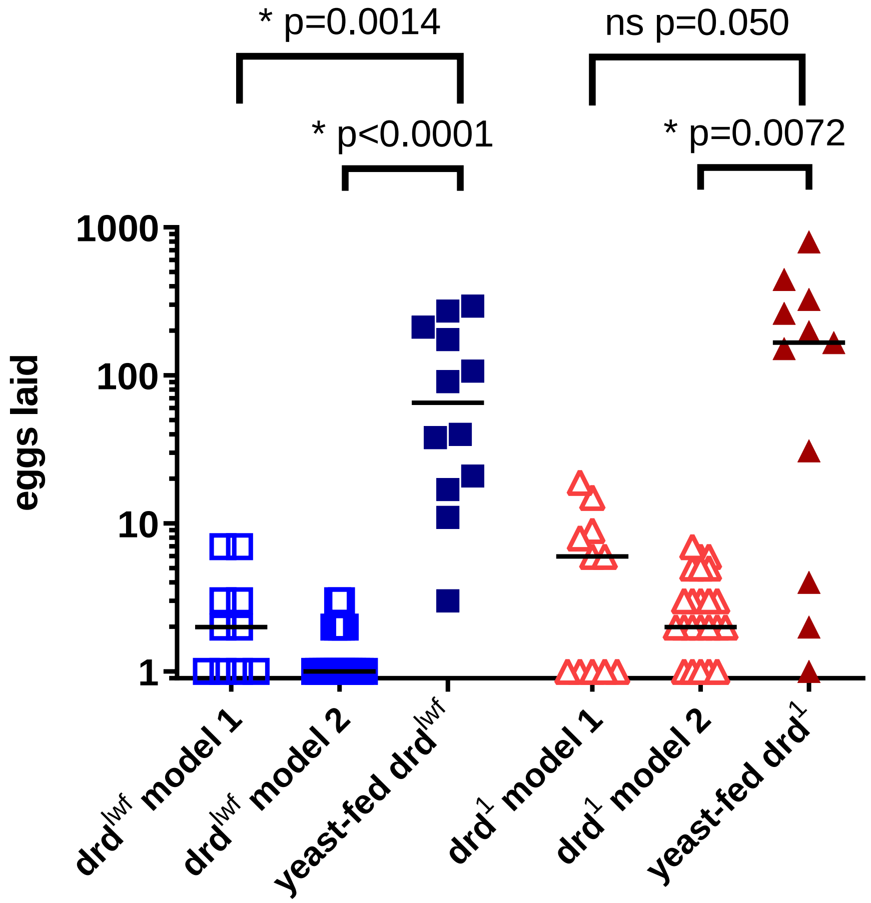

### Figure S3

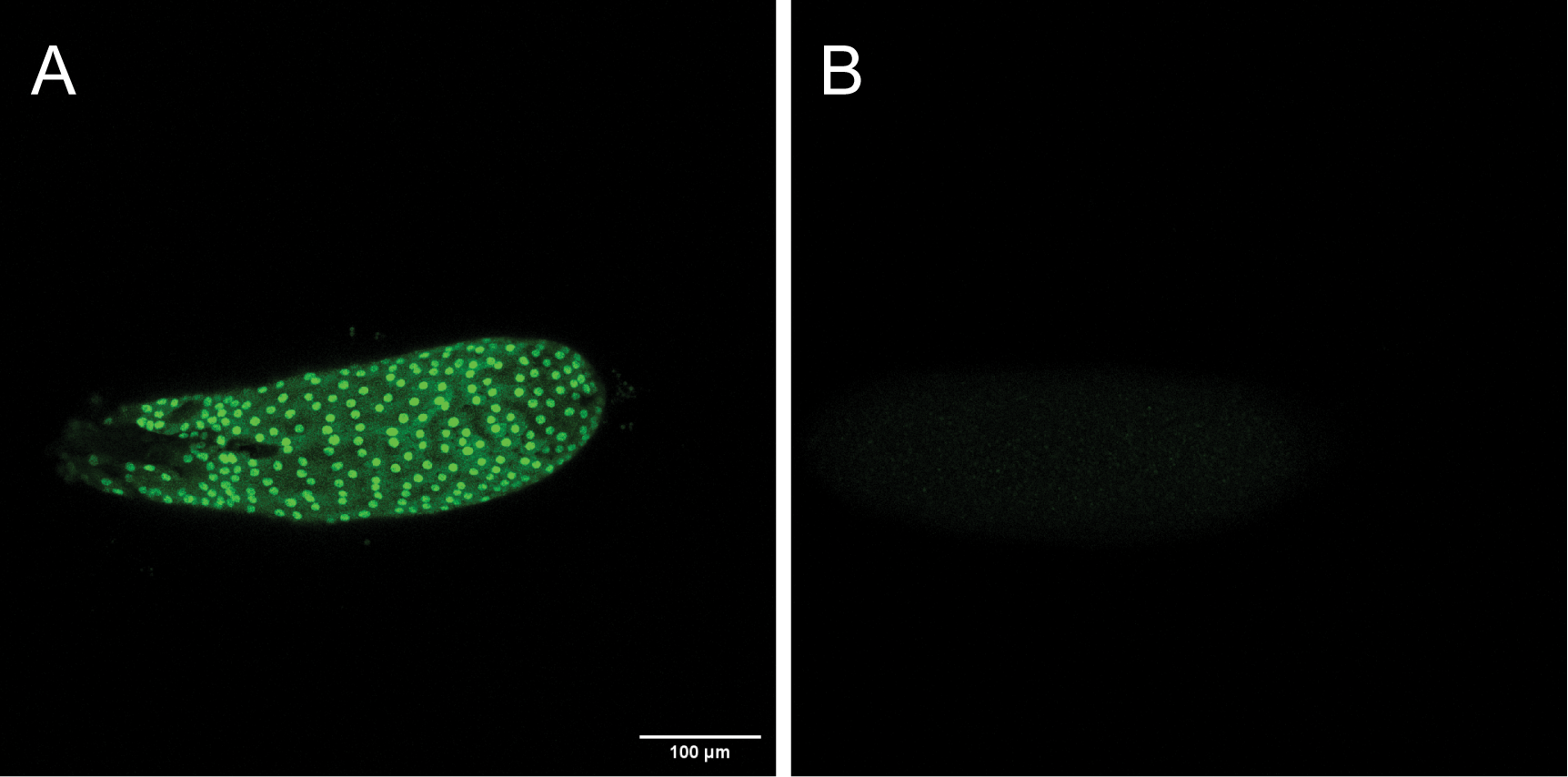
